# A phase separated organelle at the root of motile ciliopathy

**DOI:** 10.1101/213793

**Authors:** Ryan L. Huizar, Chanjae Lee, Alex A. Boulgakov, Amjad Horani, Fan Tu, Kevin Drew, Edward M. Marcotte, Steven L. Brody, John B. Wallingford

## Abstract

Hundreds of different cell types emerge in the developing embryo, each of which must compartmentalize cell type specific biochemical processes in a crowded intracellular environment. To study cell type specific compartmentalization, we examined motile ciliated cells, which must assemble vast numbers of dynein motors to drive ciliary beating, as mutation of dyneins or their assembly factors causes motile ciliopathy. We show that dyneins, their assembly factors, and chaperones all concentrate together in <u>Dyn</u>ein <u>A</u>ssembly <u>P</u>articles (DynAPs). These phase-separated organelles are specific to ciliated cells but share machinery with stress granules. Our data suggest that a common framework underlies ubiquitous and cell-type specific phase separated organelles and that one such organelle is defective in a human genetic disease.

---

Motile cilia are microtubule based cellular projections that beat in an oriented manner to generate fluid flows that are critical for development and homeostasis. Accordingly, defects in ciliary beating underlie a constellation of human diseases called the motile ciliopathies, which are characterized by *situs inversus*, chronic airway infection, and infertility (*1, 2*). Motile ciliopathies frequently result from mutations in genes encoding subunits of the multi-protein dynein motors that drive ciliary beating (Fig. 1A) (*1, 2*).

**Figure 1.**
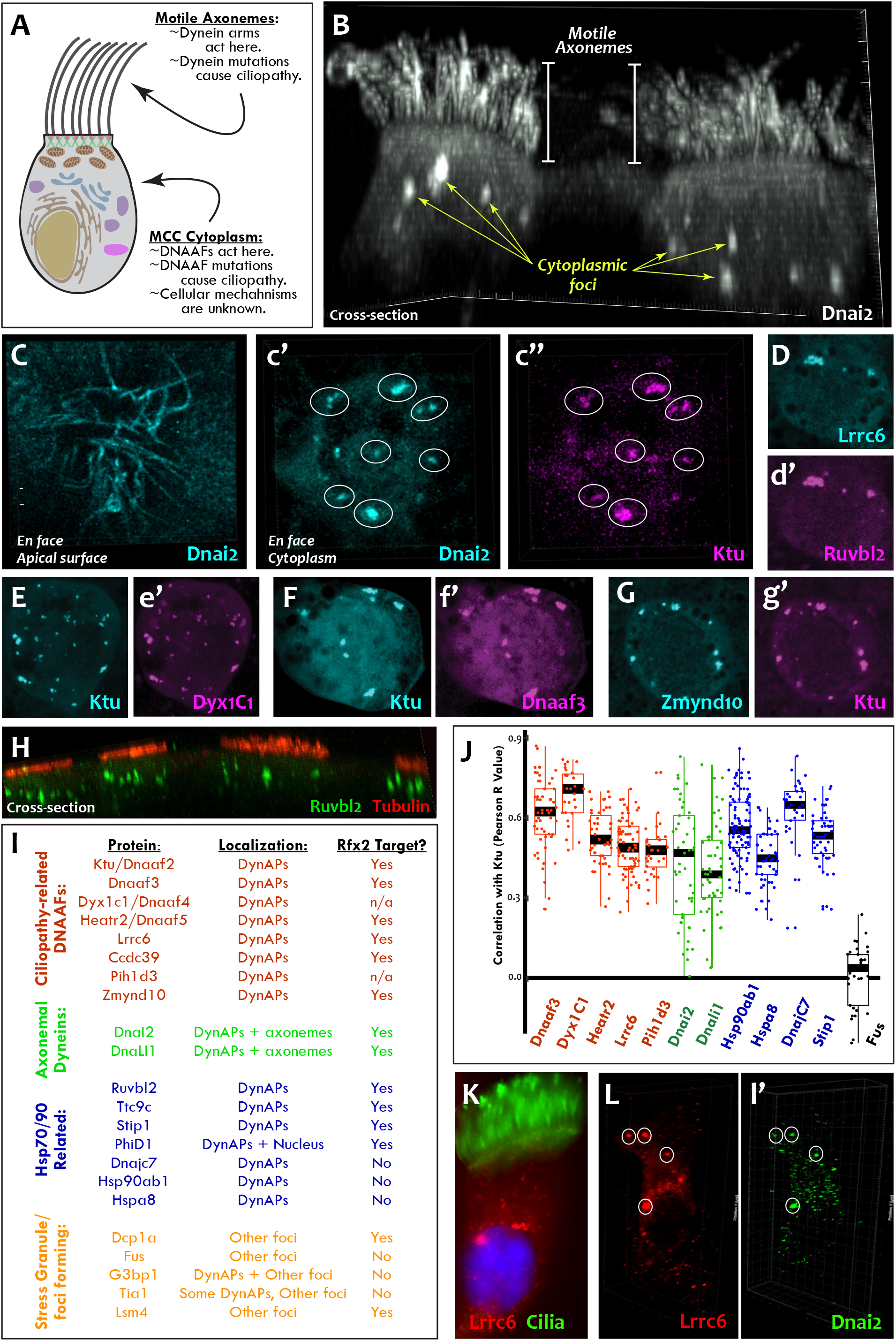
DNAAFs, Dyneins and chaperones co-localize together in DynAPs. **A.** Schematic showing a multiciliated cell indicating the site of function for proteins linked to motile ciliopathy. **B.** Cross-sectional projection of mucociliary epithelium; GFP-Dnai2 in MCCs localizes to both axonemes and cytoplasmic foci. **C.** *En face* projection showing GFP-Dnai2 localization in motile axonemes. **c’.** *En face* projection through the cytoplasm of the cell shown in C reveals GFP-Dnai2 in foci (circles). **c’’.** RFP-Ktu co-localizes in foci containing GFP-Dnai2 (circles). **D-G.** *En face* optical sections showing co-localization of FP fusions to the indicated proteins. **H.** Cross-sectional projection shows immunostaining of endogenous Ruvbl2 (green) and cilia (acetylated tubulin, red). **I.** Partial table of proteins examined in this study (see Supp. Table 1 for complete listing). **J.** Pearson correlations of pixel intensity for FP fusions to indicated proteins compared to GFP-Ktu. **K.** Immunostaining of primary human MCC reveals endogenous Lrrc6 (red) in cytoplasmic foci, cilia are marked by acetylated tubulin (green). **L.** Lrrc6 labeled foci in human MCCs (red) are enriched in Dnai2 (green in l’).

Interestingly, dynein motors are pre-assembled in the cytoplasm before being deployed to cilia (*3*), and it is now clear that many motile ciliopathies result from mutations in an array of cytoplasmic Dynein Arm Assembly Factors (DNAAFs) (Fig. 1A) (*4–16*). Previously, we reported that Heatr2/Dnaaf5 is present in cytoplasmic foci in human MCCs (*8*); and interestingly, we found that many other foci-forming proteins in MCCs are encoded by genes controlled by Rfx2 (*17*), a component of the evolutionarily conserved motile ciliogenic transcriptional circuitry (*18–22*). Because no unifying cell biological mechanism for DNAAF action has yet emerged, we set out to explore the link between the motile ciliogenic transcription factors, cytoplasmic foci, and dynein arm assembly.

## Dynein Assembly Particles (DynAPs) are evolutionarily conserved organelles compartmentalizing axonemal dyneins and their assembly factors

To probe the cell biology of the dynein assembly process, we generated fluorescent protein (FP) fusions to DNAAFs, axonemal dynein subunits, and other proteins of interest (Fig. 1I; Supp. Table 1). We assessed the localization of these proteins by confocal microscopy in the large multiciliated cells (MCCs) of *Xenopus* embryos, because these cells reliably model the biology of their mammalian counterparts, but are more amenable to *in vivo* imaging and dynamic analyses (*23*).

First, we examined two axonemal dynein subunits encoded by motile ciliopathy genes, DnaI2 and DnaLI1 (*1*). Strikingly, 3D projections revealed that in addition to the expected localization in motile axonemes, these dyneins localized strongly to cytoplasmic foci (Fig. 1B; Supp. Fig. 1A). This dual localization was also apparent in orthogonal (*en face*) projections taken from the apical cell surface (Fig. 1C) and from the cytoplasm (Fig. 1c’; Supp. Fig. 1B). We then examined eight different motile ciliopathy-associated DNAAFs (Fig. 1I, red text), and remarkably, all were localized to similar cytoplasmic foci (Fig. 1D-G; Movie 1; Supp. Fig. 1C). Likewise, the Hsp90 co-chaperone Ruvbl2 has been identified as a DNAAF in zebrafish (*24*), and Ruvbl2-GFP was present in similar foci in *Xenopus* MCCs (Fig. 1, d’). These foci were not an artifact of FP-induced aggregation (e.g. (*25*)), as immunostaining for endogenous Ruvbl2 revealed similar foci in MCCs (Fig. 1H, Supp. Fig. 1D, Movie 2).

In all cases, these foci were irregularly shaped, measured roughly 1–3 microns across, were typically present in the middle third of the apicobasal axis of MCCs, and in *en face* optical sections, tended to reside near the cell periphery (Fig. 1B-H; Supp. Fig. 1). This similar morphology prompted us to ask if these foci may represent a common compartment. By expressing pairs of differently tagged proteins, we found that all DNAAFs and dynein subunits tested co-localized extensively in foci (Fig. 1C-G, Supp. Fig. 1). This co-localization was confirmed by quantification both in 2D (Fig. 1J, red, green) and in 3D (Supp. Fig. 2).

**Figure 2.**
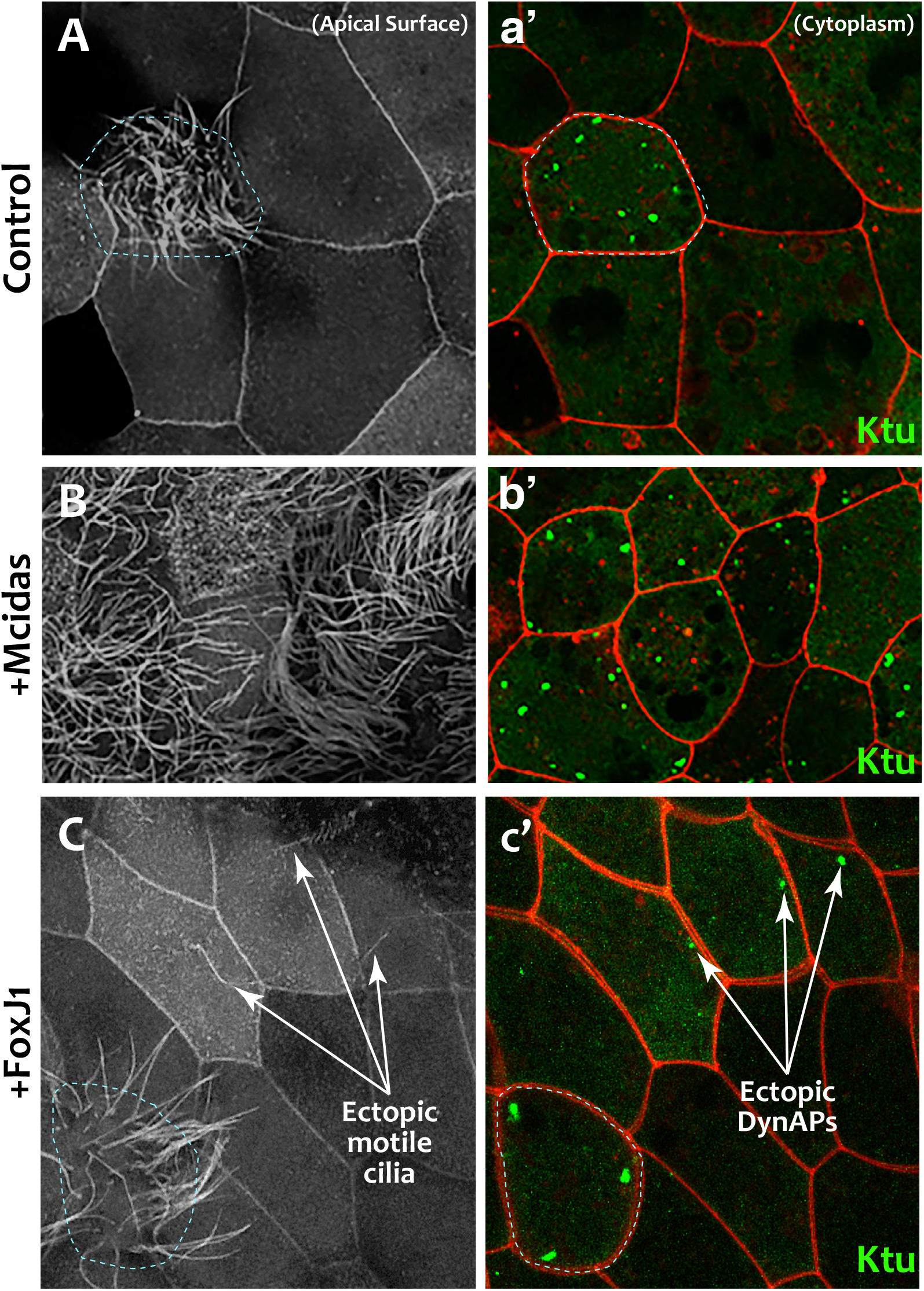
DynAPs are MCC specific and controlled by the motile ciliogenic transcriptional circuitry. **A.** Membrane labeling at the apical surface reveals a single MCC (cilia, dashed circle) surrounded by non-ciliated goblet cells. **a’.** Projection of the cytoplasm of the same cells in A; GFP-Ktu is expressed throughout, but forms foci only in the MCC. **B.** Membrane labeling at the apical surface reveals that expression of Mcidas converts all cells to MCCs. **b’.** Projection of the cytoplasm of the same cells in B; GFP-Ktu forms foci in all cells upon expression of Mcidas. **C.** Membrane labeling at the apical surface reveals that expression of Foxj1 induces solitary ectopic cilia. **c’**. Projection of the cytoplasm of the same cells in C; GFP-Ktu forms solitary foci in cells with ectopic cilia

Importantly, these DNAAF-labeled foci appeared to be novel structures, as we observed no co-localization with FP-fusions to several known foci-forming proteins, including the P body proteins Dcp1a and Lsm4 and the amyloid forming protein Fus (Fig. 1J, black; Supp. Fig. 3A-C). DNAAFs also did not co-localize with Centrin4, a basal body marker or Ccdc78, a marker of deuterosomes, dedicated organelles that generate the multiple centrioles in MCCs (*26*) (Supp. Table 1).

**Figure 3.**
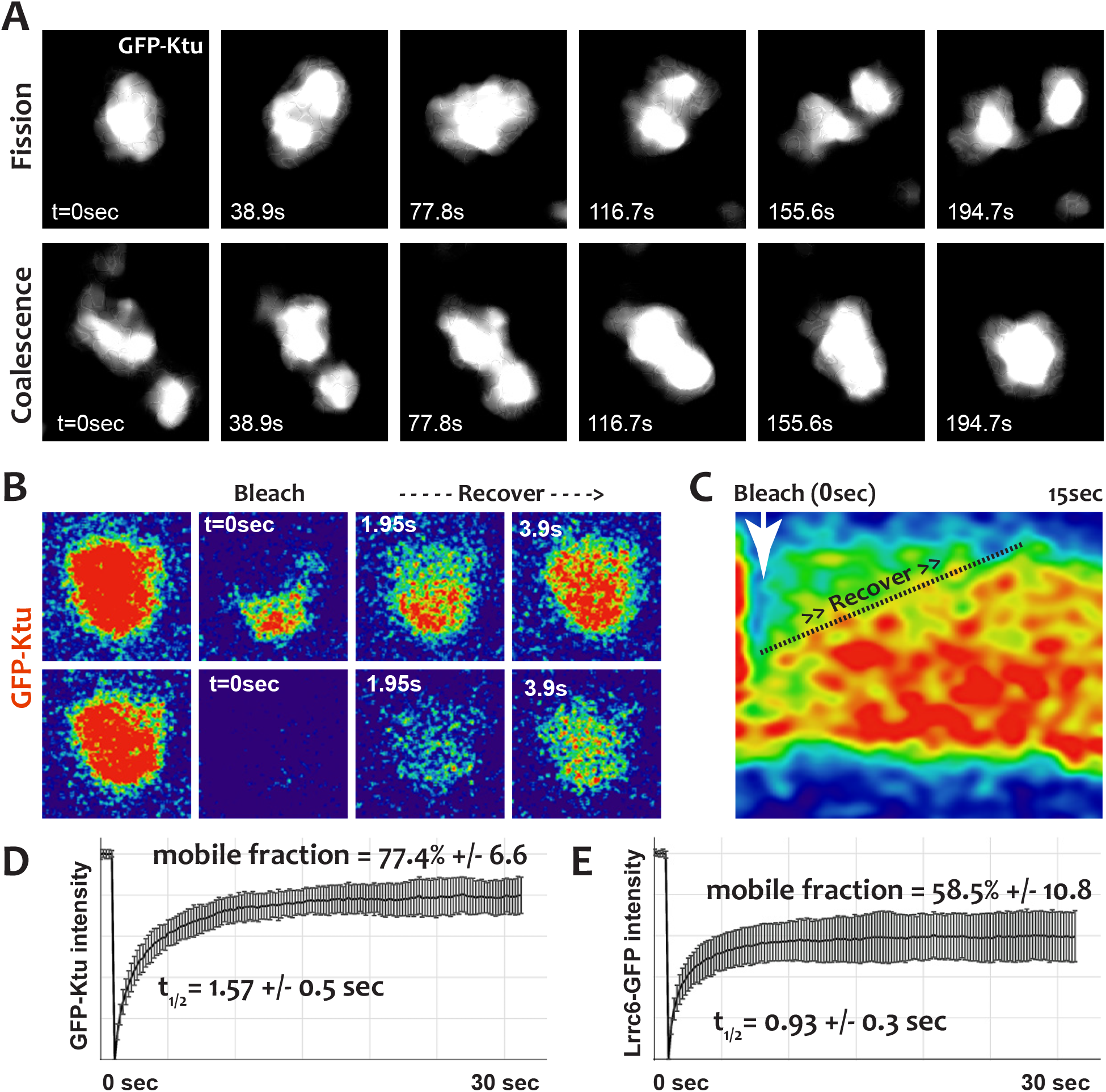
DynAPs display liquid-like behaviors. **A.** Stills from time-lapse imaging of an individual DynAP labeled with GFP-Ktu undergoing fission (upper) and later coalescence (lower) (time in seconds). **B.** Time-lapse images of GFP-Ktu recovery after photobleaching of partial DynAPs (upper) or entire DynAPs (lower). **C.** Kymograph of GFP-Ktu recovery after “half-bleach,” showing preferential intra-DynAP mobility. **D.** FRAP kinetics of GFP-Ktu after bleaching entire DynAPs. **E.** FRAP kinetics of Lrrc6-GFP after bleaching entire DynAPs.

To ask if the patterns of co-localization we observed in *Xenopus* are a conserved feature of MCCs, we re-examined primary human MCCs and found that the ciliopathy-associated DNAAF LRRC6 (*7, 10*) was present in foci that also were enriched for the axonemal dynein subunit DNAI2 (Fig. 1K, L, l’). Thus, DNAAFs and axonemal dynein subunits are specifically co-enriched in discrete cytoplasmic compartments in the MCCs of humans and *Xenopus*. We propose that these compartments represent novel organelles in which multiprotein dynein arms are assembled, and we propose the term *DynAPs*, for <u>Dyn</u>ein <u>A</u>ssembly <u>P</u>articles.

## DynAPs contain core Hsp70/90 chaperones and specific co-chaperones

Little is known about the molecular mechanisms of DNAAF function, but proteomic experiments suggest links to the Hsp70/90 chaperone machinery (*4, 14–16, 24*). Consistent with the idea of DynAPs as specialized compartments for DNAAFs, these organelles also concentrated Hsp70/90 components (Fig. 1I, blue). Core subunits of Hsp70 (Hspa8) and Hsp90 (Hsp90ab1) were strongly co-localized with DNAAFs (Fig. 1J, blue; Supp. Fig. 4A, B). Moreover, specificity of Hsp70/90 function is imparted by co-chaperones, and we identified several that co-localized strongly with DNAAFs in DynAPs (Fig. 1I, J, blue; Supp. Fig. 4C-E). Finally, we also identified Ttc9c as a DynAP component (Supp. Fig. 4F), which is of interest because the function of this protein is essentially unknown, but it binds Hsp90 and is implicated in cilia beating (*27, 28*). Thus, DynAPs represent a specialized compartment where ubiquitous Hsp70/90 machinery and specific co-chaperones are concentrated together with DNAAFs and dynein subunits.

**Figure 4.**
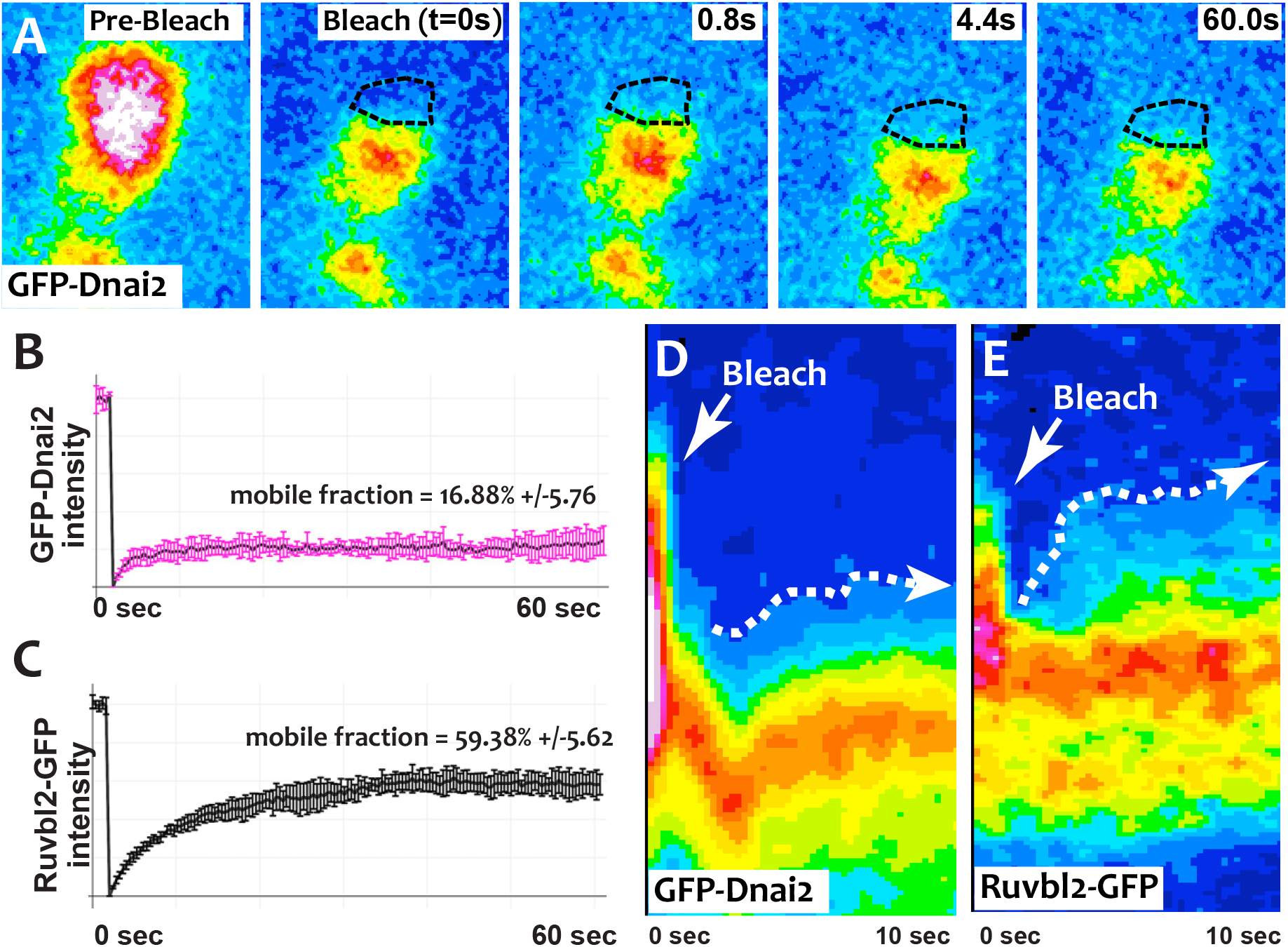
DynAP stably retain axonemal dynein subunits. **A.** Time-lapse images of GFP-Dnai2 recovery after partial photobleaching of a DynAP reveals little recovery even after 60 seconds. **B.** Frap kinetics of Dnai2 after complete bleaching of DynAPs. **C.** Frap kinetics of Ruvbl2 after bleaching of entire DynAPs. **D.** Kymograph of the first 10 seconds after bleaching shows very little recovery of GFP-Dnai2. **E.** For comparison, a similar kymograph shows more rapid intra-DynAP recover after partial bleach of Ruvbl2-GFP.

## DynAPs are MCC-specific organelles that assemble under the control of the motile ciliogenic transcriptional circuitry

We next sought to understand the developmental biology of DynAP formation. As in the mammalian airway, *Xenopus* MCCs bearing motile cilia work in concert with intermingled mucus-secreting goblet cells (*23*), but immunostaining for endogenous Ruvbl2 reported foci specifically in MCCs (Supp. Fig. 1D; Movie 2). This result suggested that DynAPs might represent MCC-specific organelles, an idea reinforced by the fact that most genes encoding DynAP-localized DNAAFs, dyneins, and chaperones are controlled by the transcription factor Rfx2, with the exception of the more broadly acting Hsp70/90 components (Fig. 1I).

To test the cell type specificity of DynAPs, we experimentally expressed FP-fusions to DNAAFs and axonemal dynein subunits throughout the mucociliary epithelium. Strikingly, even when broadly expressed, these fusions assembled into foci only in MCCs and not in the adjacent goblet cells (Fig. 2A; Supp. Fig. 1A). This MCC-specificity was not a general property of FP fusions, as the P body marker Dcp1a (*29*) assembled into foci in both MCCs and goblet cells (Supp. Fig. 5A), as did the stress granule marker G3bp1 (*29, 30*) (Supp. Fig. 5B).

**Figure 5.**
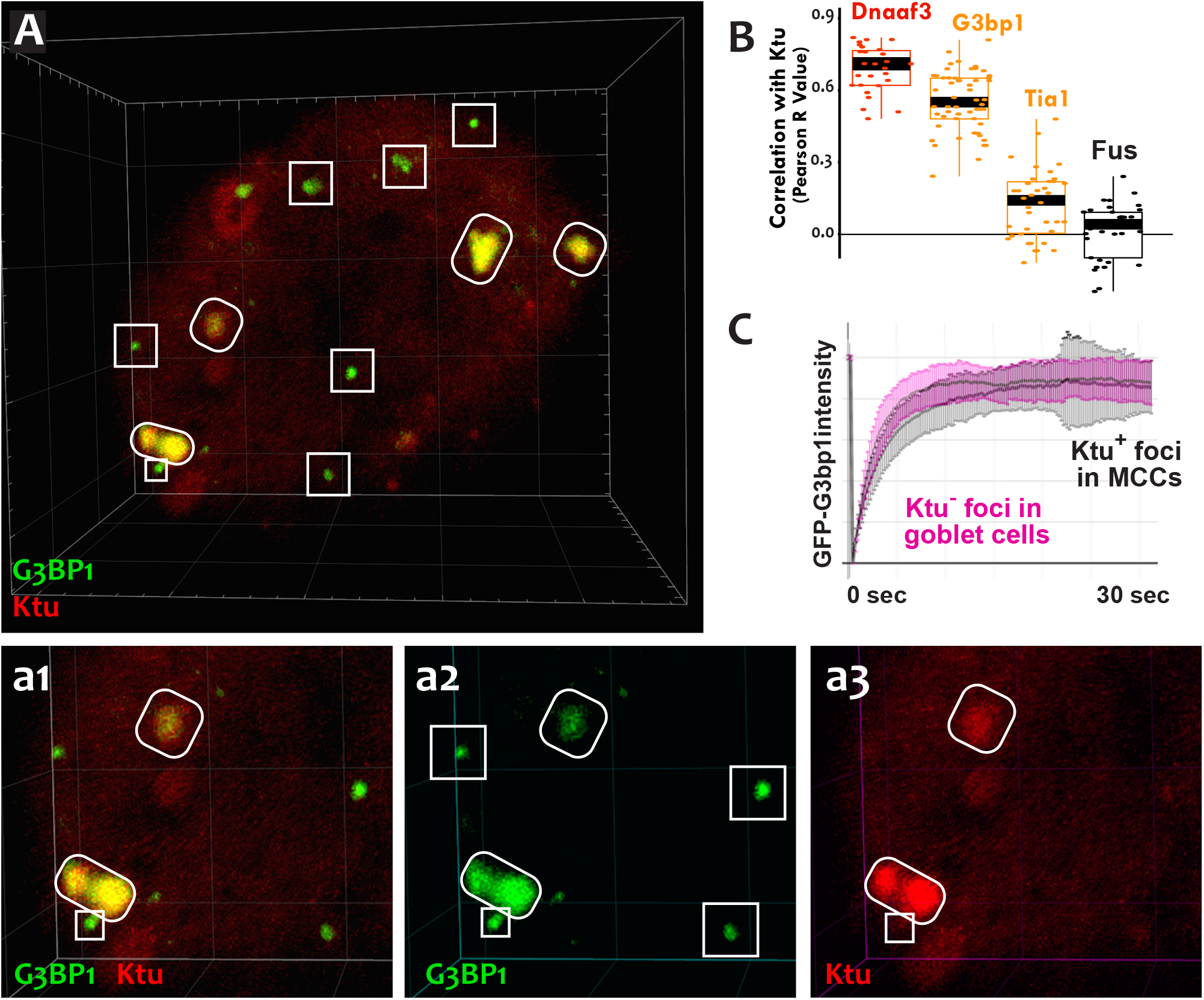
DynAPs share molecular and physical properties with stress granules. **A.** GFP-G3bp1 strongly co-localizes with DNAAFs in DynAPs (ovals), but also labels smaller foci that do not contain DNAAFs (boxes). **a1-a3.** Higher magnification views of the bottom left corner of the MCC shown in panel A. **B.** Quantification of co-localization relative to RFP-Ktu (Dnaaf4 and Fus data from Fig. 1 are recapitulated here for comparison). **C.** FRAP kinetics of GFP-G3bp1 in Ktu-positive DynAPs (black) in MCCs and in Ktu-negative foci in neighboring goblet cells (pink).

For a more direct test, we turned our attention to the motile ciliogenic transcription factors. We avoided experiments with Rfx2, as this protein acts redundantly with other Rfx family proteins; instead we examined Mcidas and Foxj1, as either is sufficient to drive motile ciliogenesis, even in normally non-ciliated cells (*19–21*). We found that conversion of non-ciliated goblet cells into MCCs by expression of the master regulator *Mcidas* (*19*) was accompanied by widespread induction of DynAPs (Fig. 2B, b’’). In addition, solitary ectopic motile cilia are induced by Foxj1 expression (*21*), and we found that these cilia were associated with assembly of ectopic DynAPs (Fig. 2C, c’). Thus, DynAPs are specific features of MCCs and their assembly is controlled by the conserved motile ciliogenic transcriptional program.

## DynAPs display hallmarks of biological phase separation

We next explored the cell biological basis of DynAP assembly. DynAPs were not labeled by a general membrane marker (Movie 1) or by markers of known membrane-bound organelles such as Golgi or endosomes (Supp. Fig. 3D-F). We hypothesized, then, that DynAPs may represent biomolecular condensates formed by liquid-liquid phase separation (*31, 32*). This possibility was exciting, because while phase separation has emerged as a widespread mechanism for compartmentalizing ubiquitous cellular processes, such as RNA processing and stress responses (*31, 32*), examples of cytoplasmic phase separated organelles with cell-type specific functions in differentiating somatic cells remain comparatively rare.

One hallmark of phase-separated organelles such as *C. elegans* p-granules and mammalian stress granules is rapid fission and fusion (*29, 33*), and time-lapse imaging of DynAPs revealed similar behaviors. DynAPs underwent fission and coalescence on the order of only a few minutes (Fig. 3A; Movies 3 & 4). A second hallmark of biological phase separation is rapid exchange of material both within organelles and between the organelles and the cytoplasm (*33, 34*). Using “half-bleach” FRAP experiments, in which only a fraction of the organelle is bleached (*33, 35–37*), we observed rapid intra-DynAP exchange of Ktu (also known as Dnaaf2 (*4*))(Fig. 3B, C). GFP-Ktu displayed similarly rapid FRAP kinetics after bleaching of entire DynAPs (Fig. 3B, D, Supp. Table 2), suggesting rapid exchange of Ktu between DynAPs and the cytoplasm. Similar turnover kinetics were also observed for another DNAAF, Lrrc6 (*7, 10*) (Fig. 3E). Finally, phase-separated organelles are predicted to scale with container size, with larger cells displaying larger condensates (*38*); and accordingly, DynAPs in the 30μm diameter *Xenopus* MCCs were substantially larger than those observed in the much smaller human MCCs (Fig. 1). Thus, DynAPs display many of the hallmarks of biological phase separation.

## DynAPs stably concentrate dynein subunits

The defining function of DNAAFs is to facilitate assembly of dynein subunits into multi-protein dynein arms prior to their transport into axonemes, so we were curious to explore the dynamics of dynein flux through DynAPs. Strikingly, unlike DNAAFs, dynein subunits were stably retained inside DynAPs (Fig. 4). In “half-bleach” FRAP experiments with DnaI2, we observed very little recovery (Fig. 4A, D, Supp. Table 2), suggesting limited mobility within DynAPs. We observed similarly limited mobility in experiments bleaching entire DynAPs (Fig. 4B), suggesting little exchange of Dnai2 with the cytoplasm, in contrast to that observed for DNAAFs. Similar results were also obtained with another axonemal dynein subunit, DnaLI1 (Supp. Table 2). These results prompted us to test other DynAP-localized proteins, and we found that the DNAAF and Hsp90 co-chaperone Ruvbl2 displayed the more rapid turnover rates similar to other DNAAFs (Fig. 4C, E). Thus, DynAPs stably concentrate dynein subunits, while assembly factors and chaperones rapidly flux through. These data are consistent with a model in which DynAPs act as “reaction crucibles” (*32*) for multi-protein dynein arm assembly.

## DynAPs share molecular machinery and protein dynamics with stress granules

Our data indicating a mix of more-fluid and more-stable components in DynAPs is of interest because such a combination has been suggested as a general principle for phase-separated organelles (*39*). Indeed, in P bodies, Dcp1a displays very rapid turnover while Dcp2 is very stably retained (*40*), and in nucleoli, the fibrillarin core is more stable and the nucleophosmin containing shell is more liquid-like (*41*). In our view, the morphology and behavior of DynAPs are most reminiscent of stress granules, which display irregular shapes and contain both more-stable and more-liquid components (*29, 39*). Moreover, the enrichment of Ruvbl2 and Hsp70/90 chaperones in DynAPs (Fig. 1I) reflects similar findings in stress granules (*39*).

We identified the stress granule component G3bp1 (*29, 30*) as a direct Rfx2 target gene (Fig. 1I), so we used this protein to examine similarities between DynAPs and stress granules. First, we observed that G3bp1 was strongly enriched in DynAPs, co-localizing strongly with DNAAFs in MCCs (Fig. 5A, a2, ovals). More interestingly, G3bp1-FP in MCCs also consistently labeled a second population of smaller foci that did not contain DNAAFs (Fig. 5A, a3, boxes). FRAP experiments revealed turnover kinetics in both population of foci that were also similar to those reported for G3bp1 in stress granules (Fig. 5C) (*29*). Thus, G3bp1 is a component of at least two populations of cytoplasmic foci in MCCs, including DynAPs.

To ask if DynAP localization was a common feature of stress granule proteins, we examined Tia1, which also functions in stress granules (*42*). Tia1-FP localized to small foci in MCCs similar to the DNAAF-negative foci labeled by G3bp1 (Supp. Fig. 5C). However, unlike G3bp1, Tia1-FP was typically not present in DynAPs, and in cases where co-localization between Tia1 and DNAAFs was observed, it was weak and diffuse (Fig. 5B; Supp. Fig. 5C, arrow). Thus, despite a similar semi-liquid-like behavior, DynAPs share only a subset of molecular components with stress granules, including G3bp1, Ruvbl2, and the Hsp70/90 chaperones (*39*).

## Motile ciliopathy associated mutations alter DNAAF flux through DynAPs

DNAAFs were first identified from human genetic studies of motile ciliopathy (*4–16*), and ultimately, we sought to link our discovery of DynAPs to their role in cilia beating. We focused on Heatr2 (aka Dnaaf5), because protein loss due to nonsense mutations in this gene causes motile ciliopathy characterized by the absence of dyneins from MCC axonemes (*8, 14*). Accordingly, knockdown of Heatr2 in *Xenopus* elicited a severe disruption of ciliary beating (Fig. 6A, B; Supp. Fig. 6A). Likewise, though MCC cilia remained morphologically normal (Fig. 6C, D), we observed a near-total loss of dynein from axonemes upon Heatr2 knockdown (Fig. 6c’, d’). This loss was not reflected by a reduction in bulk Dnai2 as assessed by western blotting (Supp. Fig. 6B). Furthermore, despite the loss from axonemes, dynein localization to DynAPs appeared largely unaffected (Fig. 6E, F), suggesting that loss of Heatr2 may impact DynAP function rather than assembly.

**Figure 6.**
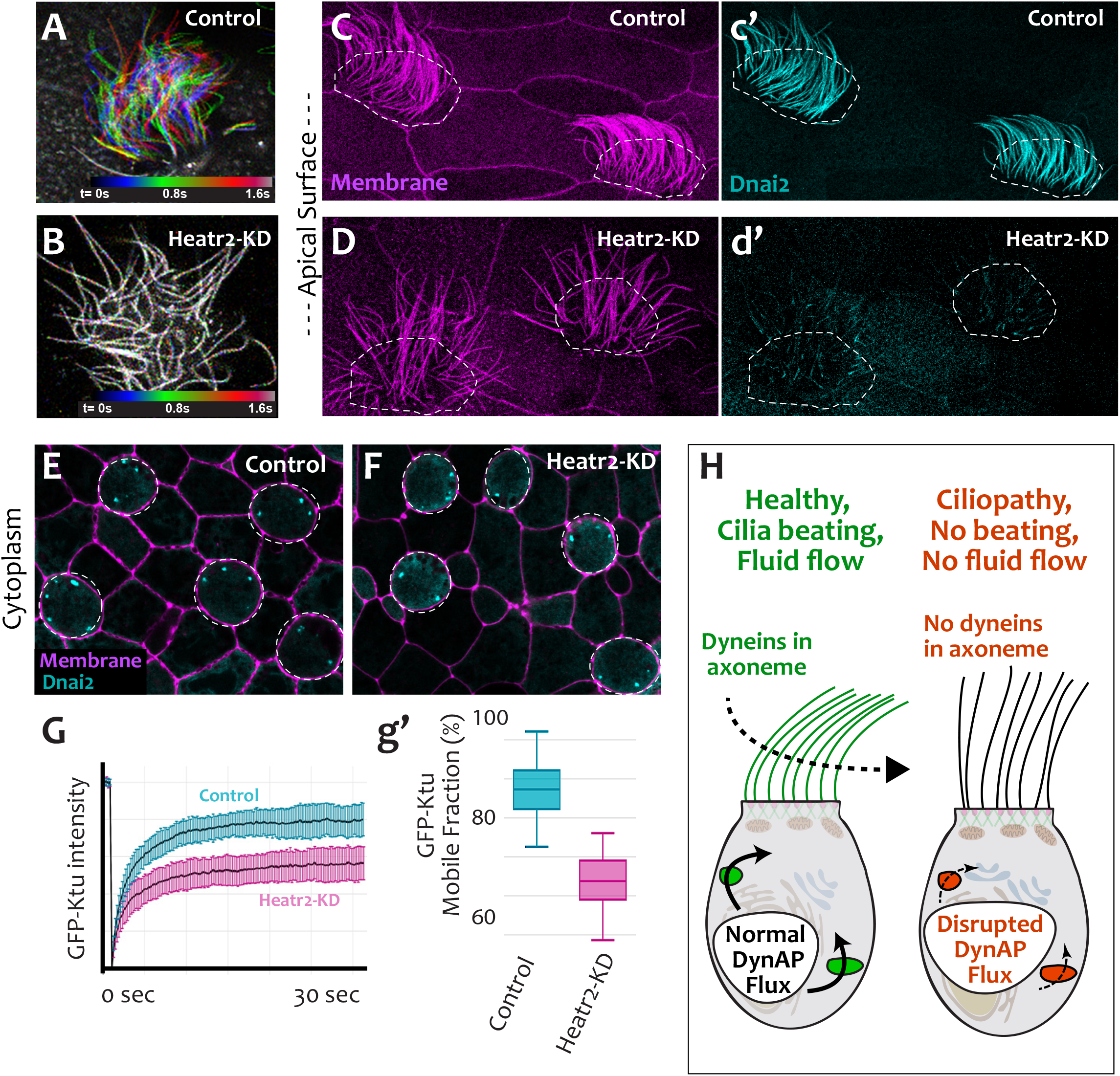
Molecular changes in human motile ciliopathy alter the liquid-like behavior of DNAAFs in DynAPs. **A.** Color-based time coding of a high-speed time-lapse movie of an MCC; Successive frames of the movie are color coded as indicated in the time key and overlaid; distinct colors in the overlay reveals ciliary movement. **B.** Similar time-coding of an MCC after Heatr2-KD; the lack of color reflects the absence of ciliary movement between frames in the movie. **C.** An apical surface view of membrane labeling (pink) reveals normal cilia morphology in control MCCs (indicated by dashed lines). **c’.** GFP-Dnai2 reveals normal localization to the motile axonemes shown in panel C. **D.** Membrane labeling of motile cilia morphology in MCCs (indicated by dashed lines) after Heatr2 KD. **d’.** GFP-Dnai2 is lost from the motile cilia shown in panel D. **E.** DynAPs are visible in an *en face* projection through the cytoplasm of MCCs (indicated by dashed lines). **F.** Despite loss from motile axonemes, Dnai2 remains localized to DynAps in the cytoplasm of MCCs (dashed lines). **G, g’.** Heatr2 knockdown significantly impairs the mobility of GFP-Ktu in DynAPs. **H.** Model for DynAP function and dysfunction in motile ciliopathy.

Consistent with this idea, FRAP analysis of superficially normal DynAPs revealed a significant reduction in the mobile fraction of GFP-Ktu after Heatr2 knockdown (Fig. 6G, g’). This result prompted us to ask if such changes in DNAAF flux may be a more general feature of motile ciliopathy, and consistent with this idea, motile ciliopathy-associated alleles of Lrrc6 (Supp. Fig. 7A)(*7*) significantly slowed the protein’s recovery half-time in FRAP experiments (Supp. Fig. 7B-C). Thus, our data suggest that ciliopathy associated molecular changes can alter DNAAF flux through DynAPs, either *in cis* (e.g. Lrrc6) or *in trans* (e.g. Heatr2 and Ktu). The altered FRAP kinetics we observe here are similar to those reported for other disease states related to biological phase separation (*35, 36*), suggesting that alteration of the liquid like behavior of DynAPs may be an important aspect of the etiology of motile ciliopathies (Fig. 6H).

## Conclusions

Multi-protein dynein arms are essential for cilia beating and their dysfunction underlies human motile ciliopathy (*1, 2*). Studies in the green algae *Chlamydomonas* revealed that dynein arms are assembled in the cytoplasm prior to their deployment to axonemes (*3*), and over the last decade, human genetic studies identified DNAAFs as causative for ciliopathy (*4–16*). Until now, however, a unifying cell biological model for DNAAF function has remained elusive. Here, we show that that DNAAFs function in DynAPs, evolutionarily conserved organelles that form specifically in MCCs under the control of the motile ciliogenic transcriptional program. DynAPs concentrate over 20 different proteins (Supp. Table 1), including dynein subunits, DNAAFs, core Hsp70/90 chaperones, specific co-chaperones, and a subset of stress granule components.

DynAPs display hallmarks of liquid-like biomolecular condensates, and our data argue that these organelles provide a privileged space in MCCs where the high local concentration of specific proteins facilitates dynein arm assembly. These findings provide a useful framework for understanding dynein assembly and motile ciliopathy, but they also provide broader insights. For example, because ciliopathy-associated changes in DNAAFs impact DynAP liquid-like behavior, this work is also significant for adding a new entry in the growing roster of human pathologies linked to defective phase separation (e.g. (*31, 32, 35, 36*)).

Finally, our description of DynAPs suggests an attractive hypothesis regarding compartmentalization of the myriad biochemical processes that arise as cell types proliferate in developing embryos. Despite the emergence of phase separation as a mechanism for compartmentalizing cellular functions, cell type-specific, phase-separated organelles remain relatively rare. Cajal bodies control genome organization in the nuclei of some cell types, and the oocyte Balbiani body facilitates these cells’ long-term dormancy (*31, 32*). But, our description of DynAP formation specifically in MCCs predicts that a wide range of cell-type specific-liquid like organelles may await discovery. Moreover, our data suggest a model whereby the molecular framework of known ubiquitous liquid-like organelles such as stress granules is differentially modified by cell type-specific transcriptional circuits in order to assemble novel organelles to achieve specific functions.

## Acknowledgments

We thank D. Dickinson for critical reading and comments on the manuscript and Z. Sun for the gift of the Ruvbl2 antibody. This work was supported by grants from the NIH (R01 HL117164; R21 GM119021, R01 HD085901 to J.B.W. and E.M.M.; and DP1 GM106408, R01 DK110520, R35 GM122480 to E.M.M. and NIH HL128370 to S.L.B.), the ATS Foundation/Primary Ciliary Dyskinesia Foundation/Kovler Family Foundation (to A.H.), NSF (to A.A.B), the Welch foundation (F-1515, to E.M.M.), and a Supplement to Promote Diversity in Health Related Research from the NICHD (to R.L.H.).

## Supplemental Figure Legends

**Fig. S1:**
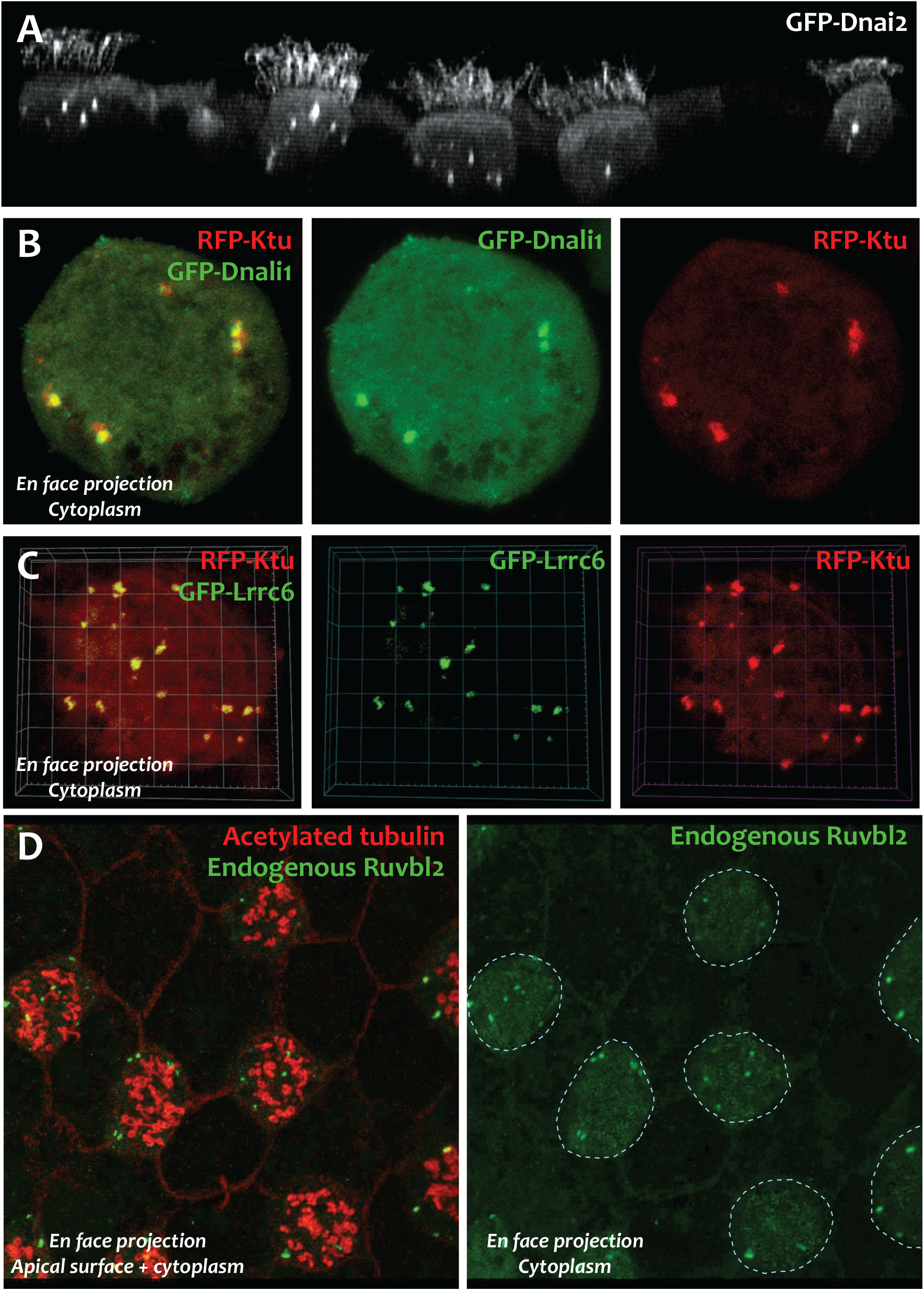
DNAAFs and Dynein subunits co-localize in DynAPs. **A.** Cross-sectional projection of a segment of mucociliary epithelium expressing GFP-Dnai2, which localizes to motile axonemes and DynAPs in MCCs. Note absence of DynAPs in the non-ciliated goblet cells adjacent to MCCs. **B.** *En face* projection through the cytoplasm of a single MCC showing GFP-Dnali1 co-localized with RFP-Ktu. Single channels of the projection are shown at right. **C.** *En face* projection through cytoplasm of a single MCC showing GFP-Lrrc6 co-localized with RFP-Ktu. Single channels of the projection are shown at right. **D.** *En face* projection through a segment of mucociliary epithelium after immunostaining for acetylated tubulin (to label cilia in MCCs, red) and endogenous Ruvbl2 (green) in DynAPs. At right is a projection through the cytoplasm of the cells shown in D; DynAPs labeled by Ruvbl2 immunostaining are present only MCCs (dashed circles).

**Fig. S2:**
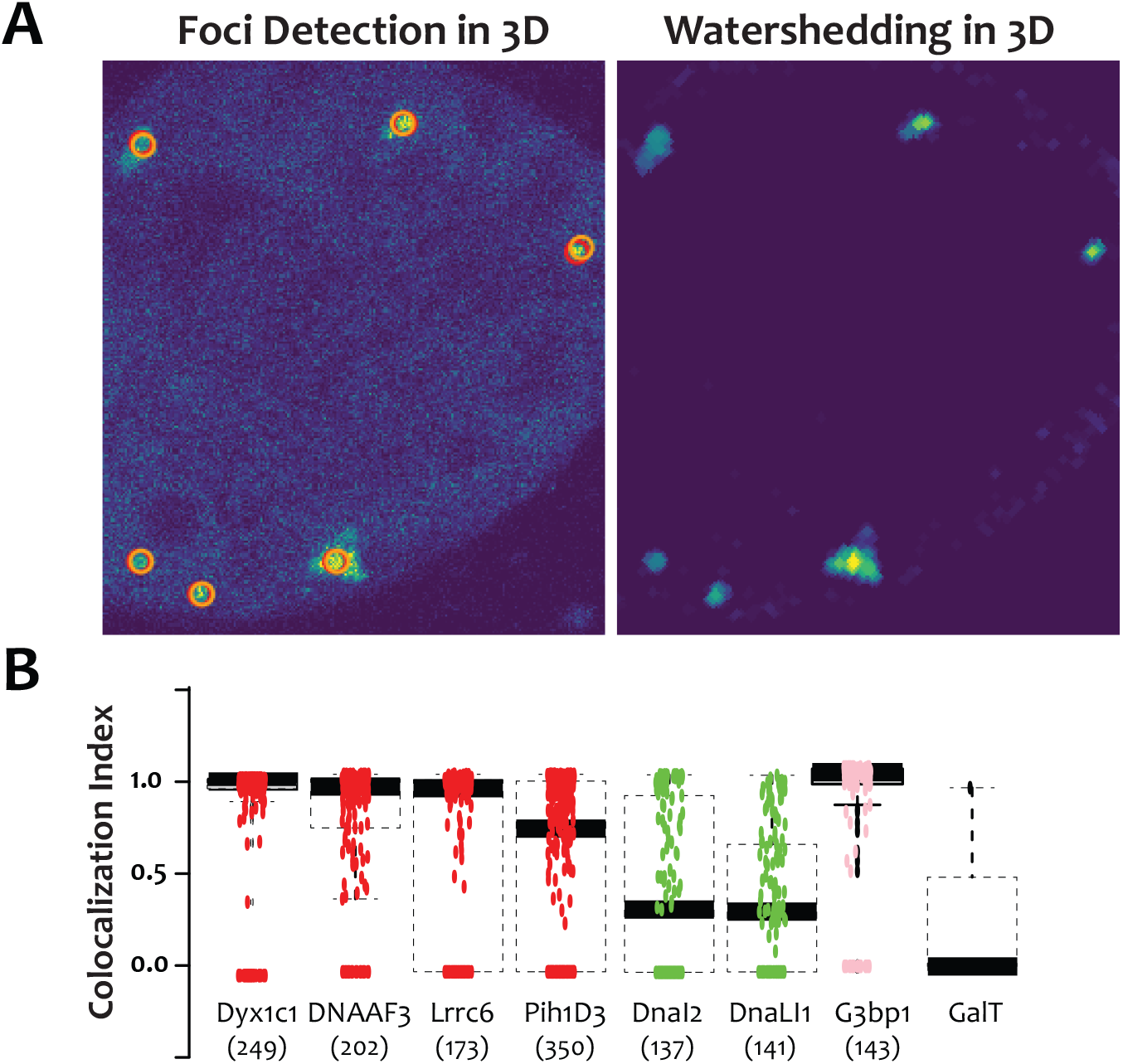
Automated 3D co-localization analysis. For unbiased assessment of co-localization, custom software was developed for automated detection of foci in confocal stacks and calculation of 3D co-localization in foci (see methods). **A.** Left panel shows focus detection in the current slice (red circle) and in the adjacent slice (orange circle) of a confocal stack. Right panel shows voxel intensity of the identified foci after watershedding. **B.** Plot of the correlation index (see methods) of FP fusions to indicated proteins relative to GFP-Ktu after automated identification of foci.

**Fig. S3:**
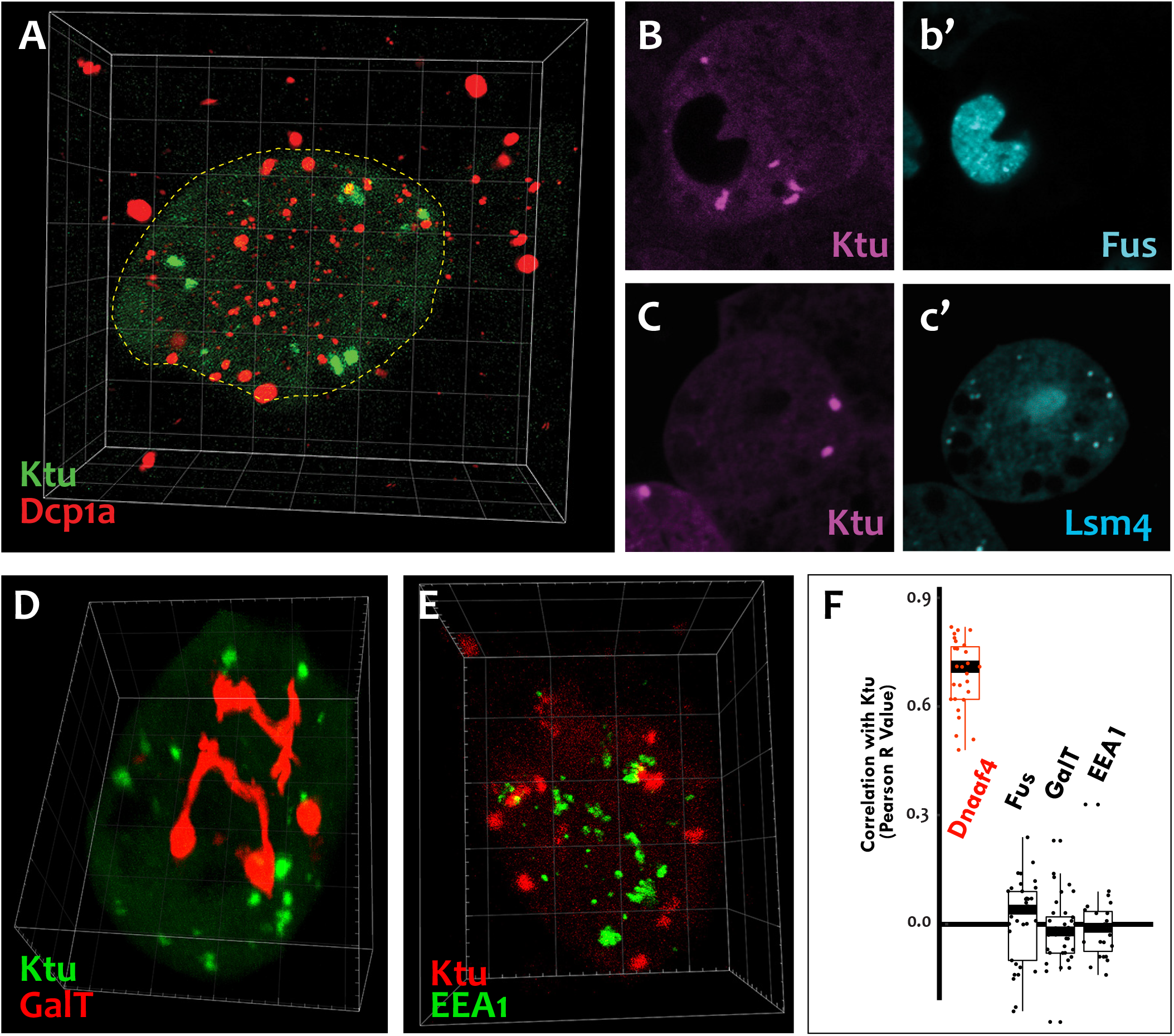
DynAPs are specific cellular compartments. **A.** GFP-Ktu labeled DynAPs (green) do not overlap with Dcp1a-labeled P bodies (red). In addition, while Dcp1a forms foci both in the MCC (dashed circle) and in the neighboring cells. **B, C.** GFP-Ktu labeled DynAps do not co-localize with foci labeled by Fus or Lsm4. **D, E.** Ktu-FP labeled DynAPs (green in D, and red in E) do not co-localize with the trans-Golgi labeled by GalT (red) or with endosome labeled with EEA1 (green). **F.** Pearson correlations of pixel intensity for FP fusions of indicated proteins compared to GFP-Ktu (For comparison, Fus and Dnaaf4 results are recapitulated here from fig. 1J).

**Fig. S4:**
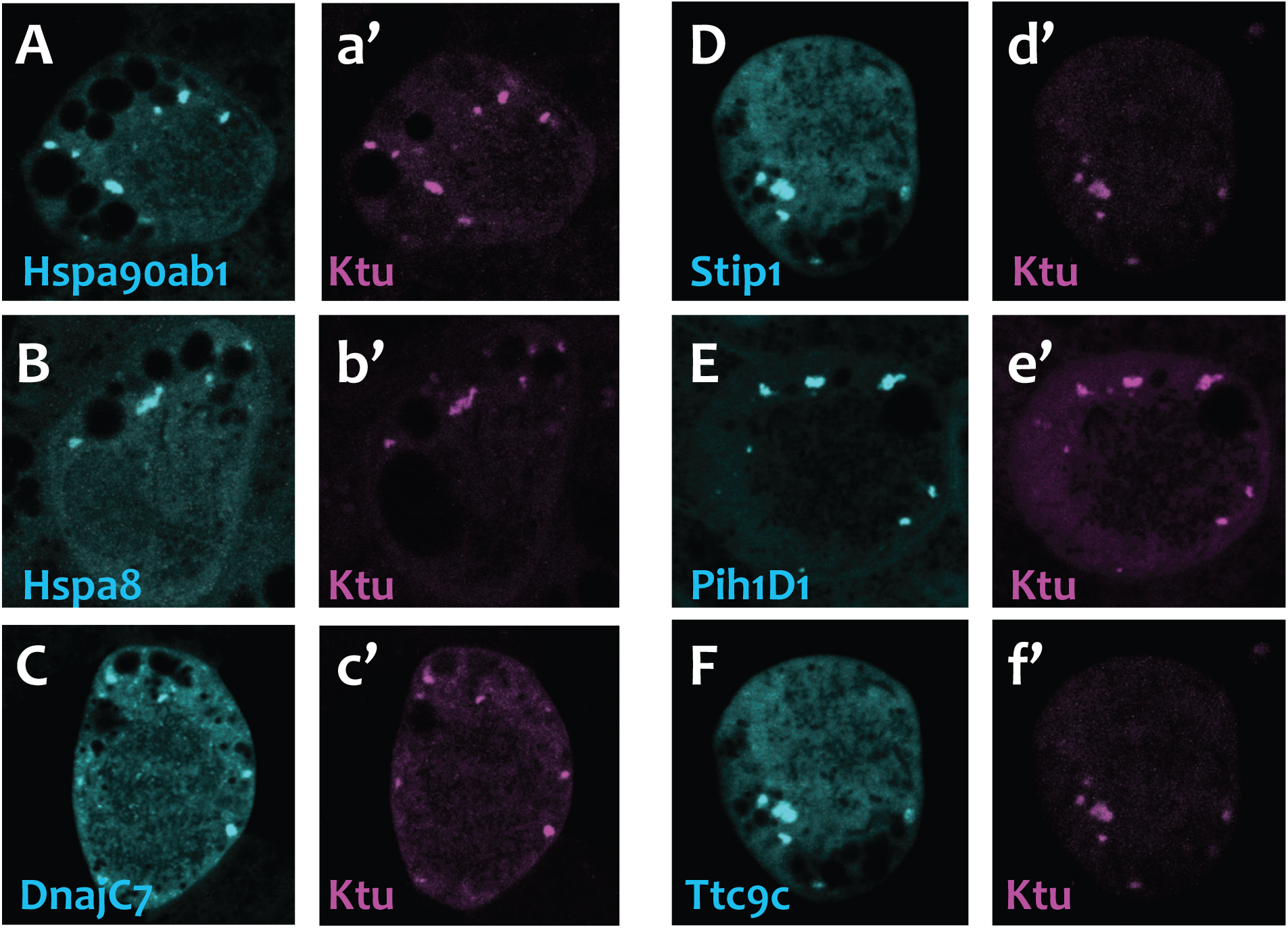
DynAPs concentrate Hsp70/90 chaperones. All left panels show GFP fusions to indicated proteins, note all co-localize in DynAPs with the RFP-Ktu shown in right panels.

**Fig. S5:**
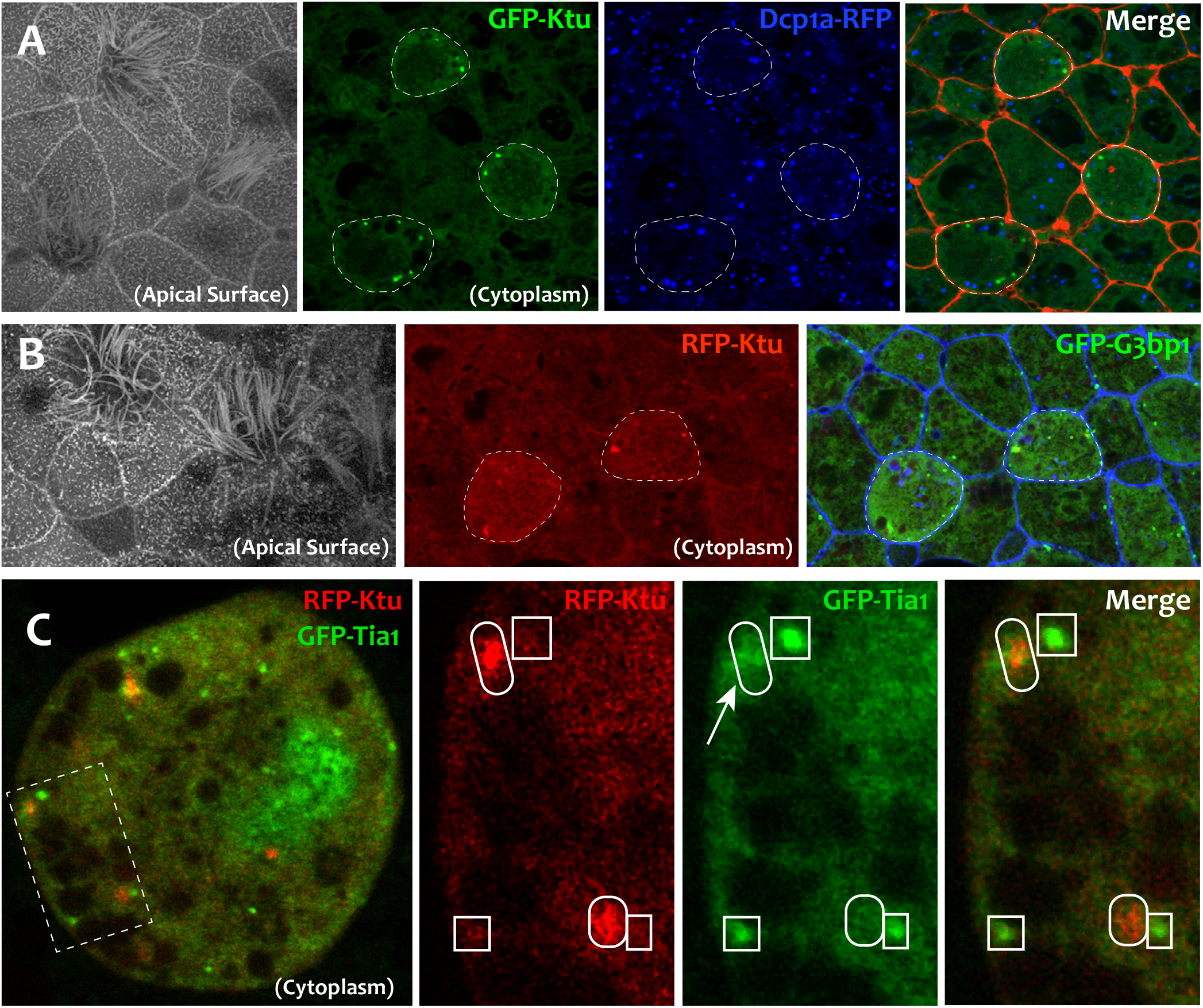
Localization of foci-forming proteins relative to DynAPs. **A.** Membrane labeling at the apical surface reveals MCCs surrounded by non-ciliated goblet cells. Panels to the right show *en face* projections through the cytoplasm of the same cells. GFP-Ktu forms foci only in MCCs (dashed circles, compare with apical surface image. Dcp1a-RFP (blue) forms foci in all cells. **B.** Membrane labeling at the apical surface reveals MCCs surrounded by non-ciliated goblet cells. Panels to the right show *en face* projections through the cytoplasm of the same cells. RFP-Ktu forms foci only in MCCs (dashed circles), while GFP-G3bp1 forms foci in all cells. Note that Ktu co-localizes with G3bp1 in DynAPs in MCCs. **C.** Projection through the cytoplasm reveals GFP-Tia1 foci (green) in a single MCC. Higher magnification views of the boxed area in B showing GFP-Tia1 (green) localizes strongly to small foci (boxes) but only weakly to the larger DynAPs (ovals) labeled by RFP-Ktu (red).

**Fig. S6:**
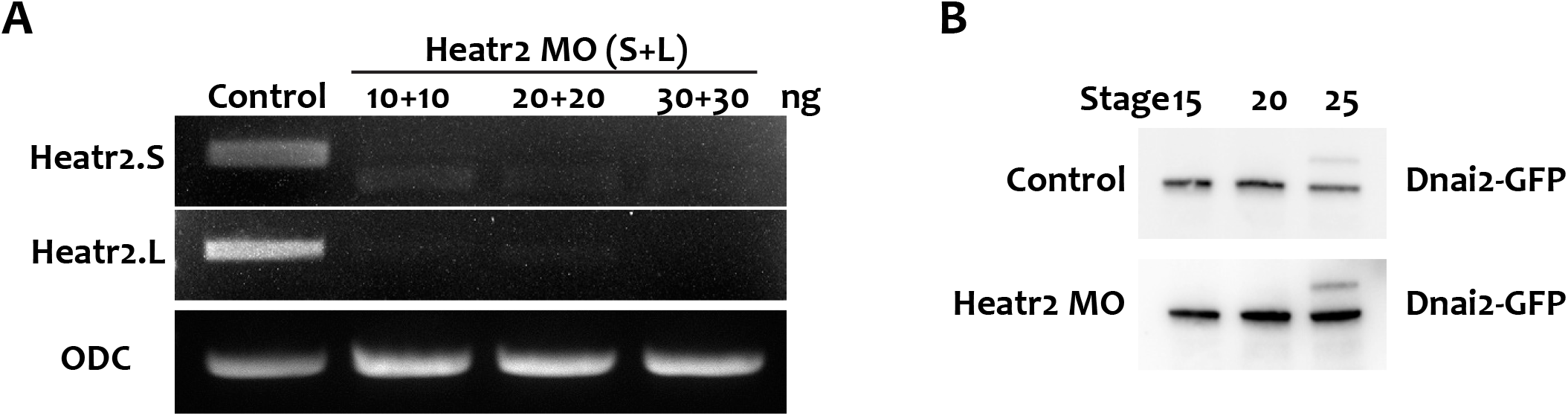
Heatr2 knockdown does not affect Dnai2 protein levels. **A.** RT-PCR shows effective disruption of splicing by injection of a combination of Heatr2-MO targeting the long and short (L&S) alloalleles of Heatr2 in the allotetraploid genome of *Xenopus laevis*. **B.** Western blotting for GFP reveals that Heatr2 KD does not impact protein levels of co-injected GFP-Dnai2.

**Fig. S7:**
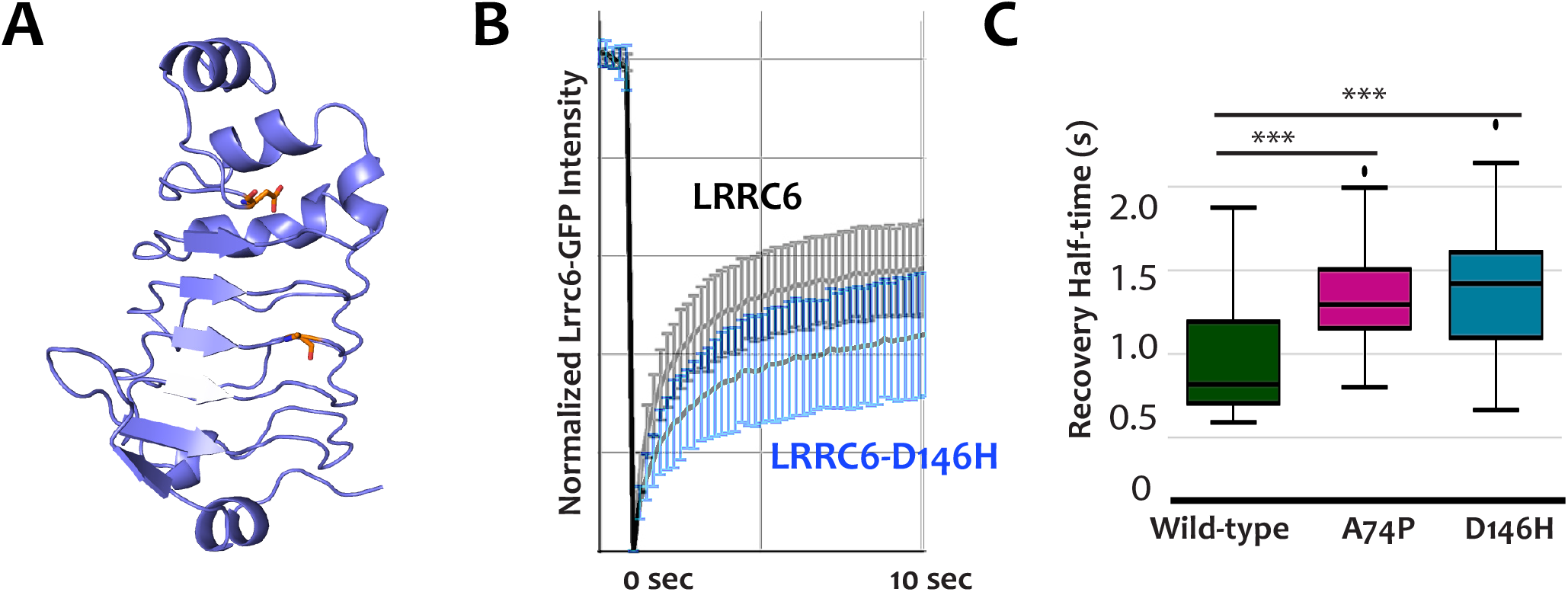
Ciliopathy-associated alleles alter recover half times of Lrrc6 in DynAPs. **A.** Structure prediction of the leucine-rich repeats of Lrrc6. Ciliopathy associated mutations (colored residues) map nearby to one another on the same external face of the repeat structure. **B.** Whole DynAP FRAP curves for GFP fusions to wild-type Lrrc6 (black) and the D146H allele (blue). **C.** The D146H and A71P alleles both slow the recovery half-time of Lrrc6.

**Supplemental Table 1:** List of proteins examined in this study. Table indicted official gene names, synonyms, functional class, overlap with KTU in co-localization studies. Rfx2 status was split into two categories: Bound indicates Rfx2 binding near the gene in ChiPseq; Regulated indicates change in expression of the gene by RNAseq after Rfx2 knockdown (*1*). Genes that are bound but not regulated likely reflect the known redundancy of cilia-related Rfx family transcription factors (e.g. Rfx3 and Rfx4).

| Gene Name | Other names | Function | Overlap with Ktu | Rfx2 status |
| --- | --- | --- | --- | --- |
| Ccdc39 |  | DNAAF | Yes | Bound, Regulated |
| Dnaaf2 | Ktu | DNAAF | n/a | Bound |
| Dnaaf3 |  | DNAAF | Yes | Not in dataset |
| Dnaaf4 | Dyx1c1 | DNAAF | Yes | Bound, Regulated |
| Dnaaf5 | Heatr2 | DNAAF | Yes | Bound |
| Lrrc6 |  | DNAAF | Yes | Bound, Regulated |
| Pih1d3 |  | DNAAF | Yes | Not in dataset |
| Zmynd10 |  | DNAAF | Yes | Bound, Regulated |
| Dnai2 |  | Dynein | Yes | Bound, Regulated |
| DnaLI1 |  | Dynein | Yes | Bound, Regulated |
| Ruvbl2 | Reptin | DNAAF | Yes | Regulated |
| DnajC7 | Ttc2 | Hsp | Yes | No |
| Hsp90ab1 |  | Hsp | Yes | No |
| HspA8 |  | Hsp | Yes | No |
| Pih1d1 |  | Hsp | Yes + Nuclear | Bound, Regulated |
| Stip1 | Hop | Hsp | Yes | Bound, Regulated |
| Ttc9c |  | Hsp | Yes | Bound, Regulated |
| Celf3 |  | Foci | No | Bound, Regulated |
| Dcp1a |  | Foci | No | Regulated |
| G3bp1 |  | Foci | Yes + other foci | Bound, Regulated |
| Lsm4 |  | Foci | No | Bound, Regulated |
| Tia1 |  | Foci | Partial + other foci + nuclear | No |
| Tial1 | Tiar | Foci | No | Bound, Regulated |
| Ccdc78 |  | Deuterosome | No | Bound, Regulated |
| Centrin4 |  | Basal bodies | No | Bound |
| Eea1 |  | Early endosome | No | No |
| Fus |  | Nuclear foci | No | No |
| Galt |  | Trans Golgi | No | Not in dataset |

**Supplemental Table 2:** FRAP kinetics for all tested proteins.

Compiled FRAP data
|  | <u>Protein</u> | <u>t1/2 (sec)</u> | <u>mobile fraction (%)</u> |
| --- | --- | --- | --- |
| Total Bleach | KTU | 1.57(0.5) | 77.4(6.6) |
|  | Lrrc6 | 0.93(0.3) | 58.5(10.8) |
|  | Ruvbl2 | 3.14(0.49) | 59.38(5.62) |
|  | Hsp90ab1 | 2.7(1.15) | 54.6(2.89) |
|  | DnaI2 | 1.0(0.33) | 16.88(5.76) |
|  | DnaLI1 | 1.13(0.65) | 22.02(5.71) |
|  | G3bp1 (DynAPs) | 2.2(0.6) | 84.1(6.4) |
|  | G3bp1 (Other foci) | 1.43(0.6) | 83.9(6.4) |
| Partial Bleach | KTU | 1.57(0.5) | 77.4(6.6) |
|  | DnaI2 | 1.04(0.66) | 20.1(9.77) |
|  | G3bp1 (DynAPs) | 1.15(0.4) | 74.8(12.2) |
|  | G3bp1 (Other foci) | 1.1(0.6) | 88.9(10.0) |
| Total Bleach, Experiments: |  |  |  |
|  | KTU + Heatr2-KD | 1.58(0.6) | 53.5(7.3) |
|  | Lrrc6-A71P | 1.36(0.3) | 61.1(6.7) |
|  | Lrrc6-D146H | 1.43(0.5) | 49(14.2) |

## Supplemental Movie Legends

**Movie 1. DynAPs form specifically in the cytoplasm of multiciliated cells.** This annotated movie shows a rotating nearest-point projection of a 3D confocal stack of a section of mucociliary epithelium labeled by expression of membrane-RFP and GFP-Ktu.

**Movie 2. Endogenous Ruvbl2 is present in DynAPs specifically in the cytoplasm of multiciliated cells.** This annotated movie shows a rotating maximum intensity projection of a 3D confocal stack of a section of mucociliary epithelium labeled by immunostaining for acetylated tubulin to label cilia (red) and Ruvbl2 (green).

**Movie 3. Dynamic behavior of DynAPs**. This non-annotated movie shows a time-lapse of Heatr2-labeled DynAPs in a single MCC.

**Movie 4. Liquid-like fission and fusion of a single DynAP**. This non-annotated movie shows smoothened data from a time-lapse movie of a single Heatr2-labeled DynAPs.

## Supplemental Experimental Procedures

### *Xenopus* embryo manipulations

Female adult *Xenopus* were induced to ovulate by injection of hCG (human chorionic gonadotropin). *In vitro* fertilization was carried out by homogenizing a small fraction of a testis in 1X Marc’s Modified Ringer’s (MMR). Embryos were dejellied in 1/3x MMR with 2.5%(w/v) cysteine at pH7.8, microinjected with mRNA or morpholinos (MOs) in 2 % ficoll (w/v) in 1/3x MMR. Injected embryos were washed with 1/3x MMR after 2hrs and were reared in 1/3x MMR until the appropriate stages.

### Plasmids, MOs and microinjections

*Xenopus* gene sequences were provided from Xenbase (www.xenbase.org) (*2*) and open reading frames (ORF) of genes were amplified from the *Xenopus* cDNA library by polymerase chain reaction (PCR). The PCR products were inserted into a pCS vector containing a fluorescence tag. The cloned genes are as follows: KTU, Heatr2, DNAI2, DNALI1, ZMYND10, LRRC6, PHI1D1, PHI1D3, Hsp90ab1, TTC9C, STIP1, DnajC7, G3BP1, Fus, and Tia1 into pCS10R-N-term GFP; DNAAF4 and Ruvbl2 into pCS10R-C-term GFP; Ruvbl2, KTU, Heatr2, DNAI2 into pCS10R-N-term mCherry; DCP1a into pCS-dest-mCherry.

Ccdc39 and Lsm4 were obtained from the Human ORFeome and DNAAF3, Dyx1c1, and EEA1 were amplified from the *Xenopus* cDNA library were cloned into pCS2+ Gateway destination vectors containing an alpha tubulin promoter and an RFP tag or a GFP tag respectively, via the Gateway LR Clonase II Enzyme. LRRC6 mutagenesis was performed using the QuickChange II Site-Directed Mutagenesis kit (Agilent Technologies).

Human GalT-GFP or RFP was derived from GalT-CFP (*3*) by exchange of CFP. Capped mRNAs were synthesized using mMESSAGE mMACHINE SP6 transcription kit (ThermoFisher Scientific). Morpholino antisense oligonucleotides (MOs) against *Heatr2* were designed to block splicing of mRNAs transcribed from both L and S alloalleles (Gene Tools). Heatr2 MO sequences and injected doses are as follow:

Heatr2 MOs : Heatr2.L (30 ng) : 5′-ACATTATCAATCACAACCTGGTATA-3′ Heatr2.S (30 ng): 5′-CATTGAATTCCTCACCTGATTTCAG-3′

For imaging, mRNAs and DNAs for fluorescence proteins were injected into two ventral blastomeres of 4 cell stage embryos with 100 pg/injection and 40 pg/injection, respectively. mRNA of FoxJ1 (*4*) was injected with 200 pg/injection into two ventral blastomeres. mRNA of MCIDAS(*5*)-hGR from CS10R-MCIDAS-hGR (100 pg/injection) was injected into all four blastomeres at the 4 cell stage and animal cap explants were dissected at stage 8, treated with 10^−7^ M dexamethasone at stage 11, cultured until stage 26 and then imaged.

### Imaging, FRAP and image analysis

Embryos expressing fluorescent proteins were fixed at stage 26 with 1x MEMFA (0.1 M MOPS, 2mM EGTA, 1 mM MgSO_4_, 3.7% formaldehyde, pH7.4) for 40 min at stage 26, washed with PBS and then imaged. For live images, *Xenopus* embryos were mounted between cover glass and submerged in 1/3x MMR at stage 25–28. Imaging was performed on a Zeiss LSM700 laser scanning confocal microscope using a plan-apochromat 63X 1.4 NA oil objective lens (Zeiss) or Nikon eclipse Ti confocal microscope with a 63×1.4 oil immersion objective. For FRAP experiments, a region of interest (ROI) was defined for full bleach experiments as a 1.75 µm × 1.75 µm box and for half-bleach experiments as a 0.8 µm × 0.4 µm box. ROIs were bleached using 50% laser power of a 488-nm laser and a 0.64 µs pixel dwell time. Fluorescence recovery was recorded at ~0.20 second intervals for up to 300 frames. Bleach correction and normalization was carried out using a custom python script (modified from http://imagej.net/Analyze_FRAP_movies_with_a_Jython_script). Frap curves were generated using Plotly. 3D projections were generated in Fiji or IMARS.

### 3D co-localization algorithm

This algorithm and the steps within was primarily implemented using scikit-image (*6*). First, to segment the cell in the image, an Otsu threshold [10.1109/TSMC.1979.4310076] is applied to the GFP channel, followed by morphological closing and opening. The largest contiguous object by area remaining in the image is the cell. Second, the Laplacian of Gaussian is used to identify puncta in both GFP and RFP channels. To reduce spurious false positives near edges, during this step only the region outside of the cell is temporarily assigned the median pixel intensity within the segmented cell. Third, background fluorescence is subtracted from the cell using a median filter. Fourth, the watershed algorithm is applied to the background-corrected image, using the identified puncta as markers. Fifth, puncta watersheds are merged across each cell’s z-stack: for each image, the set of watersheds in each image is superimposed over images directly above and below in the z-stack. Overlapping watersheds are taken to represent the same puncta in 3D space. Each channel is processed separately. This results in three-dimensional puncta watersheds distributed across each channel’s z-stack. Sixth, the watershed regions are cross-correlated between each pair of GFP and RFP images. Note that regions outside of the watershed have a value of zero, and therefore contribute nothing to the cross-correlation. For each three-dimensional puncta watershed, the individual (two-dimensional) pairwise correlations between all layers are summed to obtain the total three-dimensional cross-correlation.

### Immunostaining

*Xenopus* embryos were fixed at stage 25 by cold Dent’s fixative (80% methanol + 20 % DMSO) overnight and then were transferred in 100 % methanol. Embryos were rehydrated consecutively with TBS (155 mM NaCl, 10 mM Tris-Cl, pH7.4) and then were blocked in 10 % FBS, 5% DMSO in TBS. Monoclonal anti-acetylated alpha-tubulin antibody (T6793, sigma, 1:500 dilution) and rabbit polyclonal anti-Ruvbl2 (Abcam ab91462, 1:1000 dilution) antibody were used as primary antibodies. Primary antibodies were detected by FITC-goat anti-rabbit antibody (Sigma, 1:400 dilution) and AlexaFluor 555-anti-mouse IgG antibody (Invitrogen, 1:400 dilution).

Human airway cells were fixed and immunostained as previously described using primary and secondary antibodies.(*7, 8*) Primary antibodies and dilution used included rabbit polyclonal anti-HEATR2 (1:100, HPA020243, Sigma-Aldrich, St. Louis, MO), rabbit monoclonal LRRC6 (1:100, HPA028058, Sigma-Aldrich) and mouse mono clonal acetylated α-tubulin (1:500, clone 6-11-B1, Sigma-Aldrich). Primary antibodies were detected using fluorescently labeled, species-specific donkey antibodies (Alexa Fluor, Life Technologies, Grand Island, NY, USA). Nuclei were stained using 4′, 6-diamidino-2-phenylindole (Sigma-Aldich).

### Airway epithelial cell culture

Human airway epithelial cells were isolated from de-identified surgical excess of trachea and bronchi removed from lungs donated for transplantation. The use of these cells was exempt from human studies by the Institutional Review Board at Washington University School of Medicine. Tracheobronchial epithelial cells were expanded in culture, seeded on supported membranes (Transwell, Corning Inc., Corning, NY), and differentiated using air-liquid interface conditions as previously described and maintained in culture for up to 10 weeks.

### Structural Modeling

For fold recognition studies of Lrrc6 and Heatr2 we used pGenThreader (http://www.ncbi.nlm.nih.gov/pubmed/19429599) with default parameters searched against the latest set of PDB structures. We used the Modeller (http://www.ncbi.nlm.nih.gov/pubmed/8254673) pipeline on the pGenThreader server to build homology models based on high confident fold recognition results. Pymol (The PyMOL Molecular Graphics System, Version 1.7 Schrödinger, LLC.) was used to highlight mutations on Lrrc6 model.

### RT-PCR

To verify the efficiency of *HEATR2* MOs, both L and S MOs were injected into all cells at the 4-cell stage and total RNA was isolated using the TRIZOL reagent (Invitrogen) at stage 25. cDNA was synthesized using M-MLV Reverse Transcriptase (Invitrogen) and random hexamers. Heatr2 cDNAs were amplified by Taq polymerase (NEB) with these primers:

Heatr2.L 25F GCGACTTCCGATGTGACTAA

Heatr2.L 661R CTTCCCACTGCTGTACTGTATAA

Heatr2.S 649F GGCAATGGAAAGTCCGTAGAT

Heatr2.S 1058F CAACAACCCAGTCCGTTACA

### Immunoblotting

Embryo lysates were extracted in M-PER mammalian protein extraction reagent (Thermo Scientific), separated by SDS/PAGE in 4–20% gels (Bio-Rad) and transferred onto the nitrocellulose membrane (Thermo Scientific). The membrane was blocked overnight at 4°C in 4% nonfat dry milk in PBS containing 0.1% Tween 20 (PBST). Monoclonal mouse anti-GFP antibody (SC-9996, Santa Cruz) was used as the primary antibody and was detected by HRP-conjugated secondary anti mouse antibodies (Pierce). The membrane was incubated with SuperSignal West Femto Maximum Sensitivity Substrate (ThermoFisher Scientific) and the chemiluminescence signals were detected and visualized with an ImageQuant LAS 4000 (GE Healthcare).

